# Pan-antiviral effects of a PIKfyve inhibitor on respiratory virus infection in human nasal epithelium and mice

**DOI:** 10.1101/2023.08.11.553035

**Authors:** Jonathan Baker, Hugo Ombredane, Leah Daly, Ian Knowles, Garth Rapeport, Kazuhiro Ito

**Affiliations:** National Heart and Lung Institute, Imperial College, London, United Kingdom^a^; Pharmidex, London, United Kingdom^b^

**Keywords:** Apilimod, PIKfyve, influenza, respiratory syncytial virus, human rhinovirus, coronavirus, air-liquid interface

## Abstract

Endocytosis, or internalization through endosomes is a major cell entry mechanism used by respiratory viruses. Phosphoinositide 5-kinase (PIKfyve) is a critical enzyme for the synthesis of Phosphatidylinositol (3,5)biphosphate (PtdIns(3,5)P2), and has been implicated in virus trafficking via the endocytic pathway. In fact, antiviral effects of PIKfyve inhibitors against SARS-CoV-2 and Ebola have been reported, but there is little evidence regarding other respiratory viruses. In this study we demonstrated the antiviral effects of PIKfyve inhibitors on influenza virus and respiratory syncytial virus *in vitro* and *in vivo*. PIKfyve inhibitors, Apilimod mesylate (AM) and YM201636 concentration-dependently inhibited several influenza strains in a MDCK cell-cytopathic assay. AM also reduced the viral load and cytokine release, whilst improving the cell integrity of human nasal air liquid interface cultured epithelium infected with influenza PR8. In PR8-infected mice, AM (2mg/ml), when intranasally treated, exhibited significant reduction of viral load and inflammation and inhibited weight loss caused by influenza infection, with effects being similar to oral oseltamivir (10 mg/kg). In addition, AM demonstrated anti-viral effects in RSV A2 infected human nasal epithelium *in vitro* and mouse *in vivo*, with equivalent effect to that of ribavirin. AM also showed anti-viral effects against human rhinovirus and seasonal coronavirus *in vitro*. Thus, PIKfyve is found to be involved in influenza and RSV infection, and PIKfyve inhibitor is a promising molecule for pan-viral approach against respiratory viruses.

## INTRODUCTION

Respiratory virus infections cause an enormous disease burden both in children and in adults throughout the world. Influenza A and B viruses, respiratory syncytial virus (RSV) as well as SARS-CoV-2 are responsible for the highest numbers of deaths and hospitalizations worldwide, but other respiratory viruses, including rhinovirus, parainfluenza viruses, adenovirus, human bocavirus and seasonal coronaviruses, are also currently known to cause severe diseases in addition to mild upper respiratory tract infections, especially in patients with chronic respiratory diseases (1, 2). It is now recognized that virus induced exacerbations are preventable through COVID19 pandemic experiences (3). To tackle the respiratory virus infection, the one-target-one-drug-paradigm approach is usually taken, which is characterized by many failures and great uncertainty. Especially, the initiation of the treatment requires appropriate diagnosis of specific virus, or prophylaxis treatment is difficult without the prediction of particulate virus infection. Particularly, to prepare unforeseen pandemic or protect from several respiratory viruses, pan-viral approach is more attractive.

The endosome/lysosome system is crucial for the activity-dependent internalization of membrane proteins and contributes to the regulation of lipid level on the plasma membrane (4). Some viruses hijacked this route to deliver their genome into cells for replications (5). Phosphatidylinositol 3-phosphate 5-kinase (PIKfyve) is a phosphoinositide 5-kinase that phosphorylates phosphatidylinositol-3-phosphate (PI(3)P), synthesizing PtdIns5P and PtdIns(3,5)biphosphate, which in turn regulates endomembrane homeostasis. In fact, PIKfyve is reported to be involved in the entry of viruses into host cells, including Ebola virus, Marburg virus as well as SARS-CoV-2 (6).

Apilimod mesylate (AM) is a well-known PIKfyve inhibitor with a sub-nano molar potency, originally developed for the treatment of Crohn’s disease and rheumatoid arthritis due to inhibitory activities of interleukins 12 and 23 production (7, 8). However, the clinical trials’ results were disappointing, and the development of Apilimod for this application was halted, mainly due to poor pharmacokinetics (unexpected low plasma concentration and poor bioavailability) (9, 10). PIKfyve inhibitors have recently been re-discovered as potential anti-viral agents against filovirus and SASR-CoV-2 (6, 11–13). The inhibitor is also reported to disrupt entry or replication of RNA viruses broadly, including enterovirus, coxsackievirus B3, poliovirus, Zika virus and vesicular stomatitis virus (14). AM is reported to impair SARS-CoV-2 entry into mammalian cell lines and potently suppressed viral replication (13, 15). The results of a clinical trial against the COVID-19 (NCT04446377) are not yet available, but until improved pharmacokinetics are achieved, an issue with use possibly remains. In this report, to overcome these problems, the potential of intranasal route delivery was explored rather than oral treatment.

Historically, virus infection research was conducted using submerged culture monolayer (2D) cancer line which are susceptible to specific virus species. However, especially through the COVID19 pandemic, it was realized that positive outcomes from cancer cell line were not necessarily translational to the clinic. Air-liquid interface (ALI) cultured airway epithelium are being increasingly recognized for their ability to overcome many of the disadvantages of submerged cell culture models (16–18). It consists of pseudostratified fully differentiated cells cultured in transwell inserts, wherein the apical cells are exposed to the air and the basolateral cells submerged in culture medium, and thus a more structurally and biologically accurate representation of the human respiratory microenvironment. Attempts to repurpose existing drugs to SARS-CoV-2 treatment identified the anti-malaria drug chloroquine, which demonstrated potent anti-viral activity against SARS-CoV-2 in a submerged cancer cell culture model (Vero E6 cells) (19). However, it was later proved to have limited efficacy in clinical trials and was confirmed to have unsuccessfully reduced SARS-CoV-2 infection in ALI tissue culture models (20–22), perhaps indicating that these ALI models more accurately simulate human airway tissue responses. In addition, virus replication kinetics observed in ALI airway epithelium are similar to the outcomes from human challenge studies with SARS-CoV-2, influenza and RSV (23–26). Furthermore, we can mimic intranasal treatment using ALI two chamber systems, by applying compounds apically.

Thus, the aim of this study is to investigate the pan-viral potentials of AM using influenza and RSV infected ALI cultured airway epithelium *in vitro* and mice *in vivo* by intranasal route. The anti-viral effects of AM against human rhinovirus and seasonal coronaviruses were also evaluated *in vitro*.

## RESULTS

### Anti-viral profiles of PIKfyve inhibitors against respiratory virus panel

Firstly, the anti-viral activities of PIKfyve inhibitors Apilimod and YM201636 were assessed in *in vitro* cytopathic effect (CPE) assays in MDCK cells against influenza strains including pandemic, amantadine resistant or oseltamivir resistant. AM potently inhibited CPE induced by influenza (H1N1, H3N2, H5N1, influenza B) with IC_50_ ranged from 3.8 to 24.6 µM (Table 1, Figure 1). AM did not affect cell viability at the concentrations up to 54 µM, and consequently showed a large safety margin with respect to mammalian cell toxicity (Table 1, Figure 1). The inhibitory effects of AM were similar to or more potent than those of the known influenza anti-viral, ribavirin (Table 1). In addition, another PIKfyve inhibitor, YM201636 showed potent anti-viral effects against H1N1 A/California/07/2006, H3N2 Perth/16/2009and H5N1 Duck/MN/12525/81, suggesting a potential class effect. However, YM201636 did not inhibit influenza B at all concentrations up to 21.4 µM. As the highest concentration was found not to be toxic, further testing at higher concentrations will be required to confirm the result.

**Figure 1.**
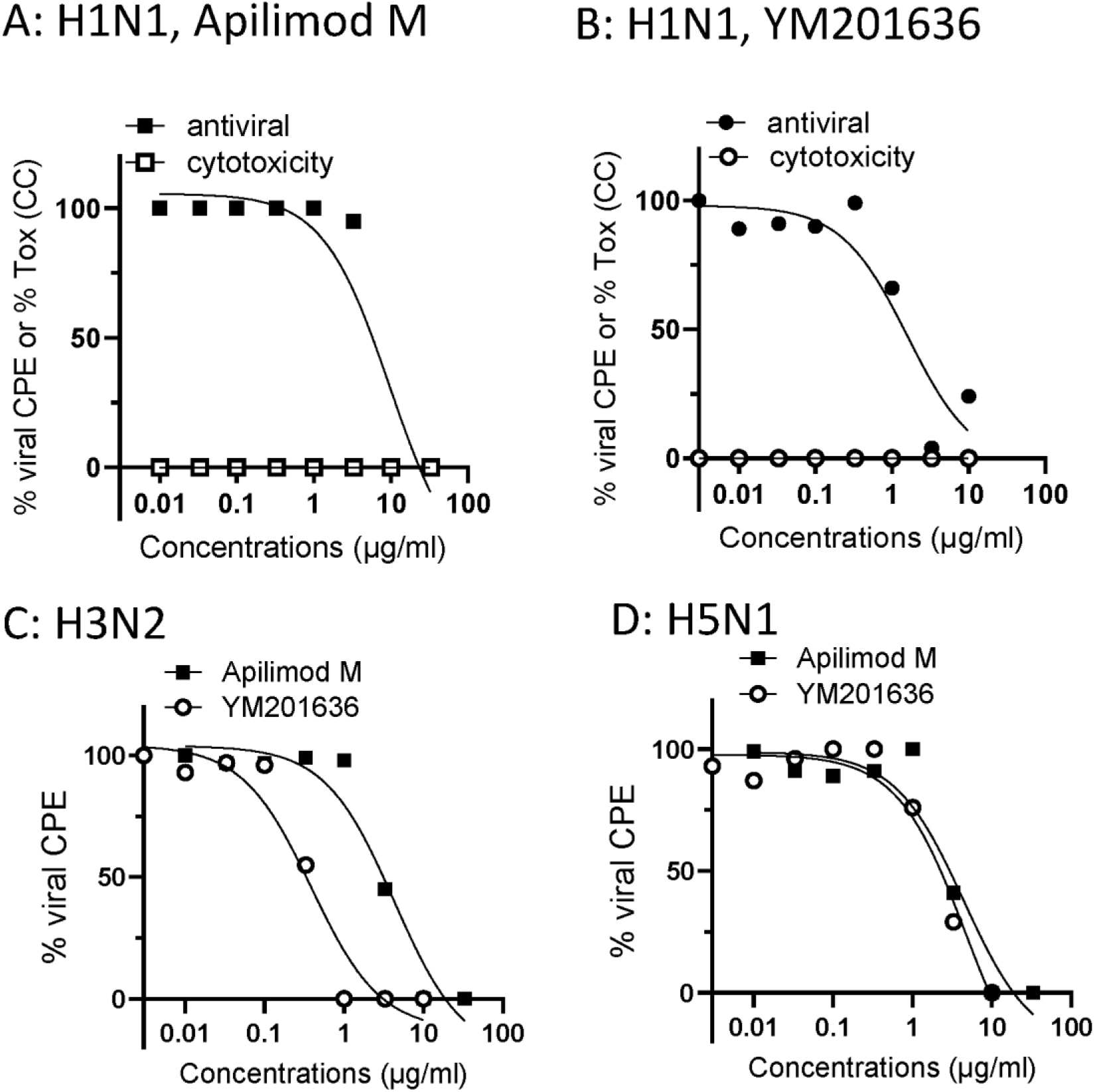
Anti-viral effects of PIKfyve inhibitors against influenza isolates. (A) Concentration-dependent cell toxicity and antiviral effects of apilimod mesylate (AM) in MDCK cells infected with H1N1 A/California/07/2009. (B) Concentration-dependent cell toxicity and antiviral effects of YM201636 (YM) in MDCK cells infected with H1N1 A/California/07/2009. (C) Antiviral effects of AM and YM in MDCK cells infected with H3N2, A/Ohio/88/2012v. (D) Antiviral effects of AM and YM in MDCK cells infected with H5N1, Duck/MN/1525/81. Compounds were treated at same time when virus inoculated, and CPE was determined Day 3 post inoculation by neutral red staining.

**Table 1.** Anti-viral effects of PIKfyve inhibitors against influenza isolates in MDCK cells, evaluated by CPE.

| Virus strain |  | Apilimod mesylate |  |  |  | YM201636 |  |  |  | Ribavirin |  |  |  |
| --- | --- | --- | --- | --- | --- | --- | --- | --- | --- | --- | --- | --- | --- |
| | | IC<sub>50</sub><br>(μM) | IC<sub>90</sub><br>(μM) | CC<sub>50</sub><br>(μM) | SI | IC<sub>50</sub><br>(μM) | IC<sub>90</sub><br>(μM) | CC<sub>50</sub><br>(μM) | SI | IC<sub>50</sub><br>(μM) | IC<sub>90</sub><br>(μM) | CC<sub>50</sub><br>(μM) | SI |
| H1N1 | A/California/07/2009 (exp1) | 9.5 | 27.8 | >54 | >5.7 | 3.0 | 21.4 | >21.4 | >7.1 | 11.1 |  |  |  |
| H1N1 | A/California/07/2009 (exp 2) | 9.2 | 31.1 | >54 | >5.9 | - | - | - | - | 25.0 | 221.1 | >4095 | >164 |
| H1N1 | A/Hong Kong/2369/2009 | 6.1 | 22.9 | >54 | >8.9 | - | - | - | - | 28.7 | 151.5 | >4095 | >143 |
| H1N1 | A/Massachusetts/15/2013 | 10.8 | 31.1 | >54 | >5.0 | - | - | - | - | 21.7 | 118.8 | >4095 | >189 |
| H1N1 | A/Louisiana/08/2013 | 6.7 | 29.5 | >54 | >8.0 | - | - | - | - | 29.5 | 176.1 | >4095 | >139 |
| H1N1 | A/Pennsylvania/30/2009 | 24.6 | 47.5 | >54 | >2.2 | - | - | - | - | 24.6 | 139.2 | >4095 | >167 |
| H1N1 | A/Michigan/45/2015 | 10.6 | 34.4 | >54 | >5.1 | - | - | - | - | 24.6 | 86.0 | >4095 | >167 |
| H3N2 | A/Ohio/88/2012v(H3N2) | 3.8 | 18.0 | >54 | >14 | - | - | - | - | 15.6 | 86.0 | >4095 | >263 |
| H3N2 | Perth/16/2009 | 5.1 | 19.6 | >54 | >11 | 0.7 | 3.4 | >21.4 | >29 | 4.9 | 32.3 | 7272 | 1480 |
| H5N1 | Duck/MN/1525/81 | 4.7 | 19.6 | >54 | >11 | 5.6 | 14.8 | >21.4 | >3.8 | 13.5 | 73.7 | 6220 | 460 |
| FluB | Florida/4/2006 | 16.4 | 47.5 | >54 | >3.3 | >21.4 | >21.4 | >21.4 | ND | 6.1 | 33.6 | 6220 | 1013 |

CPE (cytopathic effects) (1): CPE resazurin detection in 384 well format, PC786 was treated just after virus infection, and ALS-8112 was added 24hrs prior to virus inoculation, IC<sub>50</sub> IC<sub>90</sub>= concentration required for 50% and 90% inhibition, respectively, CC<sub>50</sub>= concentration to show 50% of cell death

AM and YM also exhibited potent inhibition of CPE induced by seasonal coronavirus (229E, OC43) and human rhinovirus (HRV16 and HRV1B), and particularly, the effects against seasonal CoV were very potent (Table 2). The effects of AM against RSV A2 and PIV-3 were inconclusive as it produced cell toxicity in cells used for this assay (MA104 cells, an epithelial cell line from the kidney of an African green monkey). YM did not inhibit the replication of RSV A2 and PIV3 up to 21µM (Table 2). As the concentration did not show the cell toxicity, it is inconclusive until higher concentrations are tested.

**Table 2.** Anti-viral effects of PIKfyve inhibitors against other respiratory viruses.

| Virus strain | Apilimod mesylate |  |  |  | YM201636 |  |  |  |  | Reference |  |  |  |
| --- | --- | --- | --- | --- | --- | --- | --- | --- | --- | --- | --- | --- | --- |
|  | IC<sub>50</sub><br>(μM) | IC<sub>90</sub><br>(μM) | CC<sub>50</sub><br>(μM) | SI | IC<sub>50</sub><br>(μM) | IC<sub>90</sub><br>(μM) | CC<sub>50</sub><br>(μM) | SI |  | IC<sub>50</sub><br>(μM) | IC<sub>90</sub><br>(μM) | CC<sub>50</sub><br>(μM) | SI |
| HRV14 | 12.3 | 40.9 | >54 | >4.4 | 7.5 | 23.5 | >21.4 | >2.9 | Pirodavir | 0.003 | 0.019 | 37.9 | 14000 |
| HRV1B | 0.52 | 2.8 | >54 | >103 | - | - | - | - | Pirodavir | 0.016 | 0.053 | 40.9 | 2500 |
| hCoV-229E | 0.04 | 0.23 | >54 | >1375 | 4.5 | >21.4 | >21.4 | >4.8 | M128533* | 0.19 | 1.1 | >10 | >53 |
| hCoV-OC43 | 0.007 | 0.030 | 1.3 | 193 | 0.9 | 4.1 | >21.4 | >25 | M128533* | 0.12 | 1.1 | >10 | >83 |
| RSV A2 | 19.6 | 44.2 | 29.5 | 1.5 | >21.4 | >21.4 | >21.4 | - | Ribavirin | 8.6 | 53.2 | 2047 | 238 |
| PIV-3 | 31.1 | 52.4 | 18.0 | 0.58 | >21.4 | >21.4 | >21.4 | - | Ribavirin | 73.7 | 356 | 606 | 8.2 |

CPE (cytopathic effects) (1): CPE resazurin detection in 384 well format, PC786 was treated just after virus infection, and ALS-8112 was added 24hrs prior to virus inoculation, IC<sub>50</sub> IC<sub>90</sub>= concentration required for 50% and 90% inhibition, respectively, CC<sub>50</sub>= concentration to show 50% of cell death, \*μg/ml

### Effects of Apilimod mesylate (AM) on H1N1 infection in air liquid interface (ALI)-cultured human nasal epithelial cells

The anti-viral effects of AM were also evaluated using ALI-cultured fully differentiated human primary nasal epithelium. Following inoculation with a low level of influenza PR8 strain (0.02 MOI), the quantity of virus in apical washes, as determined by TCID_50_ assay, increased from Day 1 to a peak at Day 1-2 followed by modest reduction on Day 3 (Figure 2A). We also found that the viral load waned gradually and modestly up to Day 7 (Data not shown). Treatment with AM to the apical surface, once daily from Day 0 (10 min before and 60 min together with virus inoculum, and 15 min exposure on Day 1 only), induced a concentration-dependent inhibition of H1N1 virus release on Day 2 post virus inoculation, and the effect of 0.2 mg/mL AM was statistically significant (Figure 2B). In a separate experiment, we found the effects of AM 2 mg/ml were similar to that with Oseltamivir carboxylate 0.2 mg/mL (Table 3).

**Figure 2.**
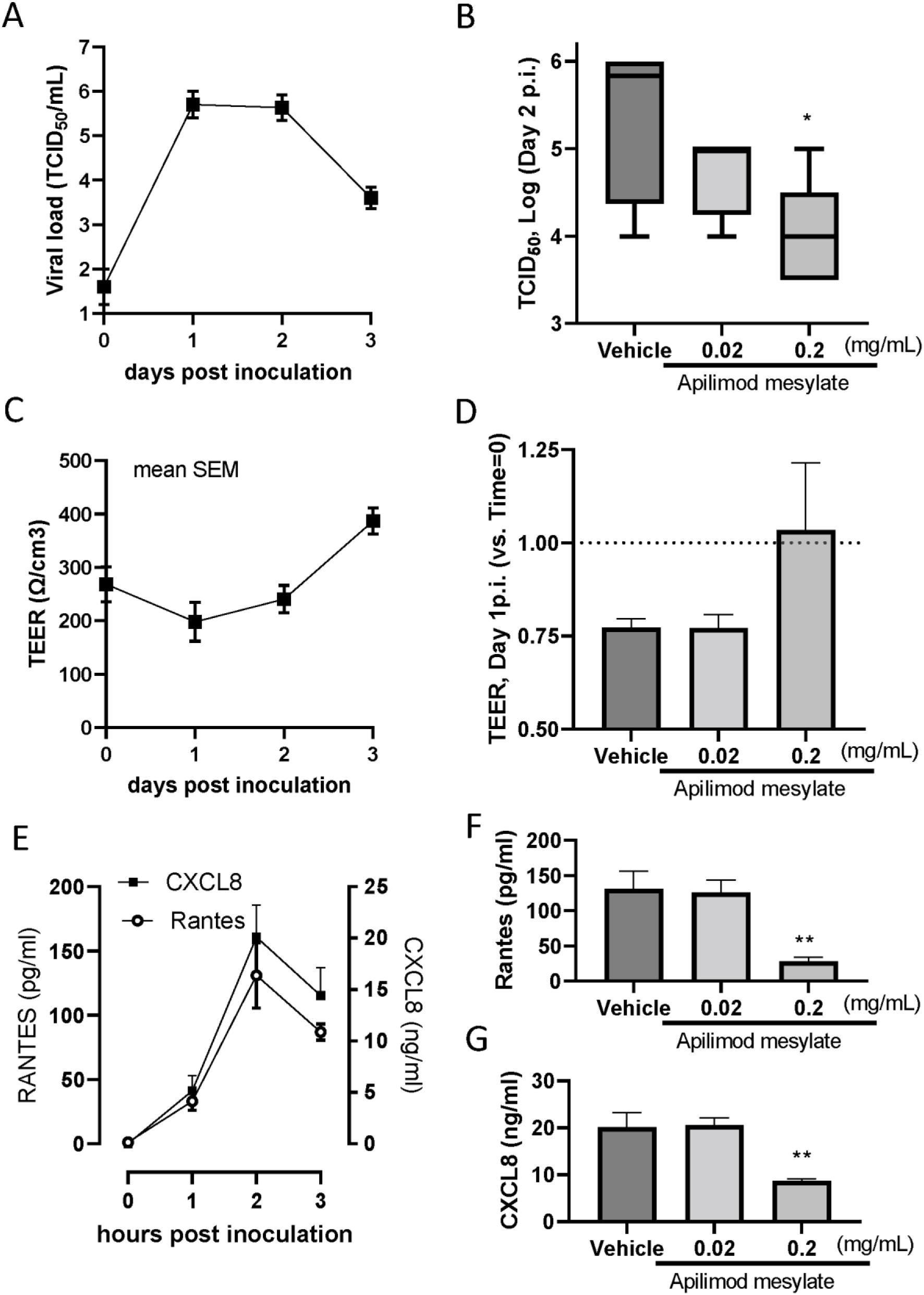
Effects of Apilimod mesylate on H1N1(PR8) infected air -liquid-interface (ALI) cultured nasal epithelium. (A) Kinetics of H1N1 viral load in apical washes post virus inoculation. On Day 0, virus was inoculated and removed after 60min absorption. Apical washes were collected post 1hr incubation (and wash) on Day 0, and once daily from Day 1 to Day 3. (B) viral load determined by TCID50 assay in apical washes from the inserts treated vehicle and AM, which were collected on Day 2 post inoculation. (C) Kinetics of cell integrity determined as TEER post virus inoculation. load in apical washes post virus inoculation. (D) TEER determined in epithelium inserts treated vehicle and AM on Day 2 post inoculation. (E) The kinetics of Rantes and CXCL8 release in apical washed from the epithelium inserts infected with H1N1, and the effects of AM on Rantes (F) and CXCL8 (G) in apical washes collected on Day 2 post inoculation.

**Table 3.** Viral load in apical wash from ALI nasal epithelium treated with Apilimod or Oseltamivir.

|  |  | Hour post H1N1 inoculation<br>(pfu/mL: geometric mean +/- SD, n=3) |  |
| --- | --- | --- | --- |
|  |  | 24h | 48 h |
| Vehicle (H<sub>2</sub>O) | apical | 2.9 ± 0.22 | 3.5 ± 0.39 |
| Apilimod mesylate<br>(0.2 mg/mL) | apical | 2.7 ± 0.30 | 2.9 ± 0.32 |
| Apilimod mesylate<br>(2 mg/mL) | apical | 2.2 ± 0.16* | 2.0 ± 0.14* |
| Oseltamivir carboxylate<br>(0.2 mg/mL) | basal | 2.1 ± 0.10* | 2.1 ± 0.20* |

\*p<0.05 (Ordinary Anova)

H1N1 infection modestly reduced cell integrity determined as TEER (trans epithelial electrical conduct) on Day1 post inoculation, and then gradually increased (Figure 2C). The level of TEER was also restored by apical AM treatment at 0.2 mg/mL although it is not statistically significant (Figure 2D). H1N1 infection also caused RANTES and CXCL8 production in apical surface, determined in apical washes. Both peaked on Day 2 post virus inoculation (Figure 2E). AM 0.2mg/ml statistically significantly inhibited the release of RANTES and CXCL8 on Day 2 post inoculation (Figure 2 F and G).

### Effects of Apilimod mesylate on H1N1 infection in mice *in vivo*

Once daily treatment with Apilimod base (AB: 0.5, 2 mg/mL) in PBS with 10% DMSO, on days 0 to 3, by intranasal (i.n.) administration, was found to inhibit body weight loss by PR8 influenza virus infection in BALB/c mice where oral treatment oseltamivir 10 mg/kg demonstrated a marked protection (Figure 3A). The effects of AM dissolved in water at 2 mg/mL (40 µg/mouse (approx. 1.6 mg/kg)) showed similar protective effects against body weight loss (Figure 3B). Viral loads in the lungs of H1N1 (PR8) infected mice was also significantly inhibited by intranasal AB and AM (both 2 mg/ml) as well as oral oseltamivir (Figure 3C). In addition, AB at 2 mg/ml showed significant reduction in the viral loads in the nasal tissue although AM showed a trend of reduction (p=0.0501) (Figure 3D). The treatment also showed the inhibition of neutrophil and lymphocyte accumulation in nasal and lung lavage on 5 days post virus inoculation (Figure 3E, Table 4).

**Figure 3.**
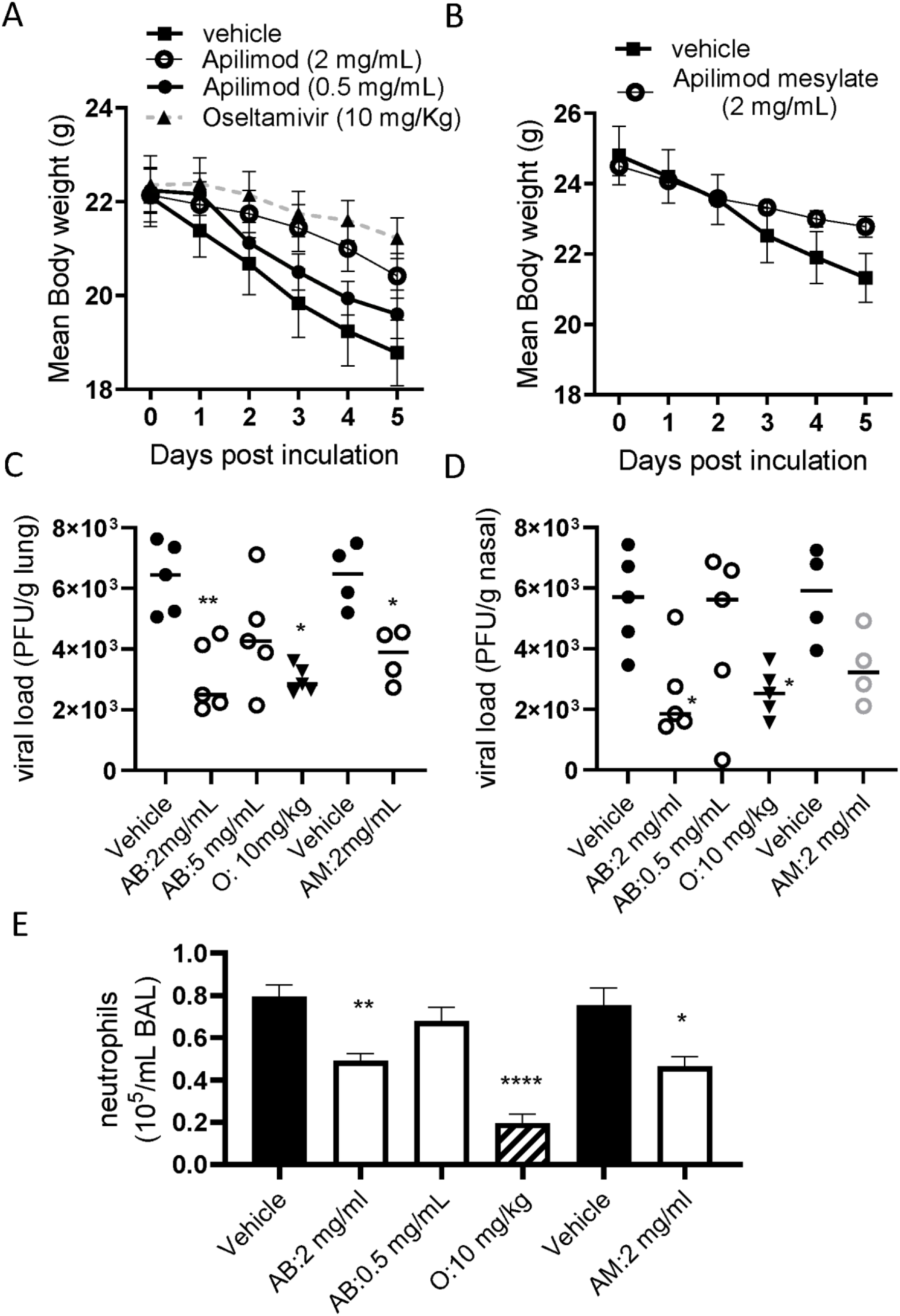
Effects of intranasal Apilimod (Apilimod base: AB, and Apilimod mesylate: AM) treatment on H1N1 infected BALB/c mice. Mice were inoculated intranasally with H1N2 PR8 (1.0 x 10^5^ PFU/mouse), and the animals were sacrificed 5 days post virus inoculation. Apilimod was treated once daily on day 0 (4 hrs before infection), and then on days 1, 2 and 3. Body weight loss triggered by virus infection (A, B), viral load in lung tissue (C) and nasal tissue (D), and neutrophil accumulation in BALF (E) were evaluated. For viral load, individual data have been plotted and Geomean. For body weight and neutrophils, mean ± SEM were shown. *\*p<0.05, ***p<0.001* vs. H1N1 infected control.

**Table 4.** Effects of Apilimod mesylate administered intranasally on H1N1 induced inflammation detected in bronchoalveolar lavage and nasal lavage of H1N1-infected mice.

| Treatment | N | Cell number (mean ± SD, $\times 10^4$, [%inhibition]) | | | Cell number (mean ± SD, [% inhibition]) | | |
| --- | --- | --- | --- | --- | --- | --- | --- |
|  |  | Bronchoalveolar lavage |  |  | Nasal lavage |  |  |
|  |  | neutrophils | Alveolar macrophage | Lymphocyte | neutrophils | Alveolar macrophage | Lymphocyte |
| Vehicle (1% DMSO saline) + virus | 5 | 7.9 ± 1.2 | 36 ± 7.0 | 7.3 ± 0.94 | 2.2 ± 0.48 | 5.1 ± 0.49 | 1.8 ± 0.14 |
| AB (0.5 mg/ml, in) + virus | 5 | 6.8 ± 1.5<br>[14] | 36 ± 4.7<br>[0] | 6.1 ± 0.42<br>[16] | 2.0 ± 0.48<br>[9] | 4.4 ± 0.35<br>[14] | 2.0 ± 0.30<br>[-11] |
| AB (2 mg/ml, in) + virus | 5 | 4.9 ± 0.76<br>[38] | 26 ± 3.3<br>[28] | 5.4 ± 0.87<br>[26] | 1.2 ± 0.38<br>[45] | 4.0 ± 0.56<br>[22] | 1.3 ± 0.10<br>[28] |
| Oseltamivir (10 mg/kg, po) | 5 | 2.0 ± 0.96<br>[75] | 11 ± 4.3<br>[69] | 2.0 ± 0.71<br>[73] | 1.2 ± 0.38<br>[45] | 3.8 ± 0.53<br>[25] | 1.3 ± 0.30<br>[28] |
| Vehicle (water) + virus | 4 | 7.5 ± 1.6 | 38 ± 3.9 | 7.5 ± 1.3 | 2.1 ± 0.57 | 4.7 ± 1.1 | 1.8 ± 0.18 |
| AM (2 mg/ml, in) + virus | 4 | 4.7 ± 0.91<br>[37] | 25 ± 4.8<br>[34] | 5.1 ± 1.1<br>[32] | 1.1 ± 0.38<br>[48] | 3.9 ± 0.76<br>[17] | 1.1 ± 0.18<br>[39] |
| Vehicle (water) + virus | 4 | 8.3 ± 2.5 | 40 ± 3.6 | 8.1 ± 1.6 | 2.2 ± 0.58 | 5.2 ± 1.0 | 1.8 ± 0.32 |
| AM (2 mg/ml, in) + virus | 4 | 6.2 ± 0.86<br>[25] | 24 ± 4.8<br>[40] | 5.0 ± 0.66<br>[38] | 1.0 ± 0.23<br>[55] | 3.6 ± 0.38<br>[31] | 0.93 ± 0.21<br>[48] |

AB: Apilimod base, AM: Apilimod mesylate, BAL: broncho alveolar lavage, NL: nasal lavage,

### Effects of Apilimod mesylate on RSV A2 infection in ALI-cultured human nasal epithelium *in vitro* and in mice *in vivo*

The anti-viral effect of AM was also evaluated using ALI-cultured fully differentiated human primary nasal epithelium infected with RSV A2. Following inoculation with a low level of RSV A2 (0.01 MOI), the quantity of RSV in apical washes, as determined by plaque assay, increased from Day 1 to Day 2 (Figure 4A). Treatment with AM to the apical surface, once daily from Day 0 (15min before virus inoculation + 60 min with virus) to Day 1 (30min incubation), induced a concentration-dependent inhibition of RSV A2 replication. Particularly, the effect at a concentration of 2 mg/ml (approximately 1.5 Log reduction) was statistically significant, which was similar to the effects of ribavirin at 100 µg/ml, treated in basal chamber, allowing exposing for 48hrs.

**Figure 4.**
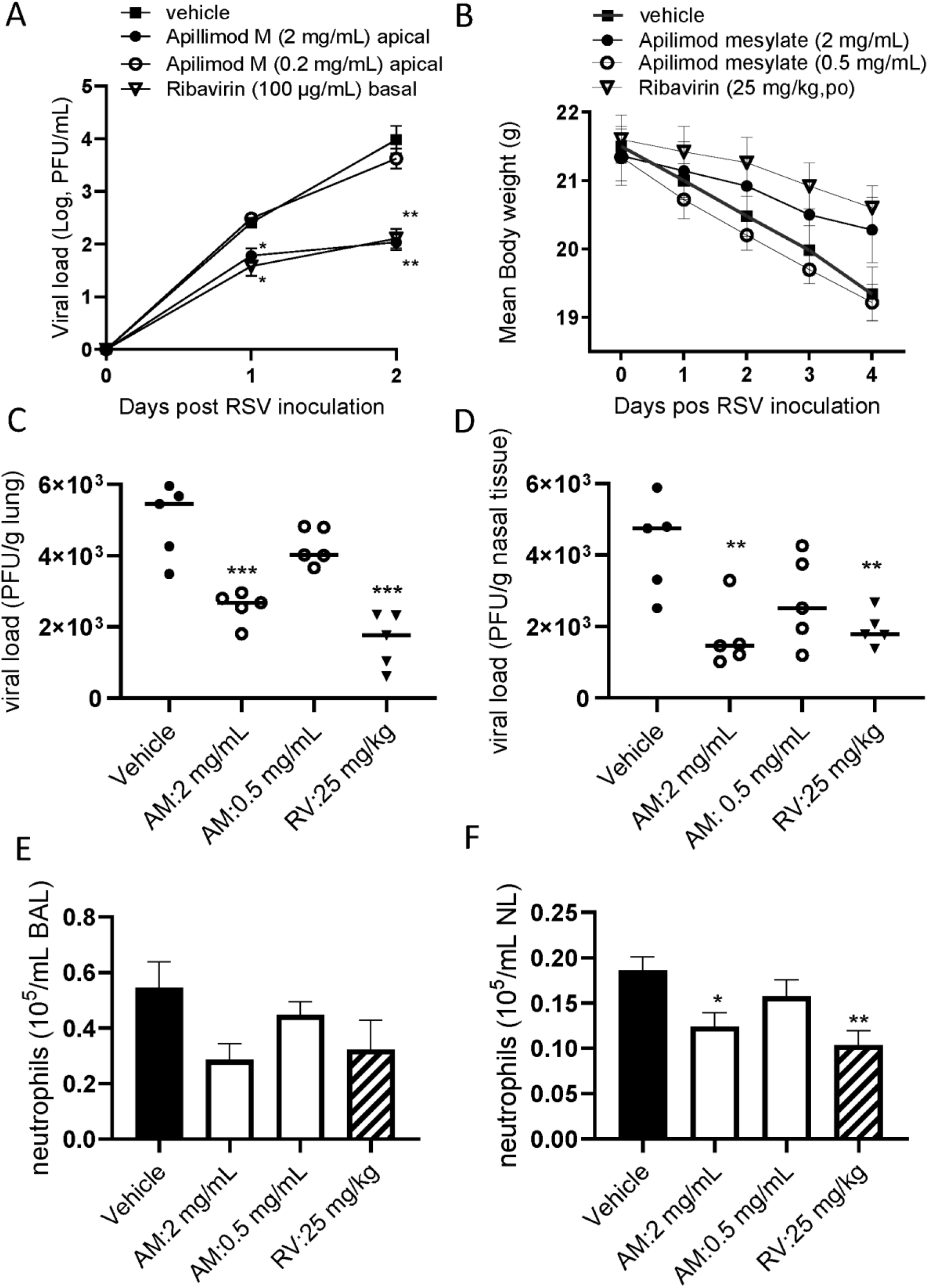
*In vitro* and *in vivo* antiviral effects of Apilimod against RSV infection. (A) Effects of Apilimod mesylate and Ribavirin on viral load in apical washes collected from H1N1(PR8) infected air -liquid-interface (ALI) cultured nasal epithelium. Apilimod was treated apically on Day 0 and Day 1 (and removed each time) or ribavirin was treated basally without removal. Viral load was determined by plaque assay (B) Effects of intranasal Apilimod (Apilimod mesylate: AM) treatment on H1N1 infected BALB/c mice. Mice were inoculated intranasally with RSV A2 (1.0 x 10^5^ PFU/mouse), and the animals were sacrificed 4 days post virus inoculation. Apilimod was treated once daily on day 0 (4 hrs before infection), and then on days 1 and 2. Body weight loss triggered by virus infection (B), viral load in lung tissue (C) and nasal tissue (D), and neutrophil accumulation in BALF (E) and nasal lavage (F) were evaluated. For viral load, individual data have been plotted and Geomean. For body weight and neutrophils, mean ± SEM were shown. *\*p<0.05, ***p<0.001* vs. RSV infected control.

Twice daily treatment with AM in mice, on days 0 to 3, by i.n. administration, was found to show beneficial effects on body weight loss triggered by virus infection where oral ribavirin at 25 mg/kg showed good protection (Figure 4B). The viral loads in the lungs and nasal tissue of RSV A2 infected BALB/c mice were significantly inhibited by 2 mg/mL intranasal AM, effects were similar to that of oral ribavirin at 25 mg/kg (Figure 4 C and D). The AM treatment also inhibited neutrophil accumulation in bronchoalveolar and nasal lavage, but the effects of AM was statistically significant only in nasal lavage.

## DISCUSSION

Considerable efforts to identify the antiviral agents to recent pandemic COVID19 were made since it was outbroken, and further works are continued to develop pan-viral agent to prepare next potential pandemic. In this manuscript, we demonstrated that Apilimod is one of promising pan anti-viral agents to tackle several respiratory virus infections.

Firstly, Apilimod (base or mesylate [water soluble form]) inhibited replication of several influenza strains including Oseltamivir/Amantadine resistant isolates in MDCK cells *in vitro* (Table 1 and Figure 1). These data suggest that Apilimod potentially inhibits influenza species broadly. YM, another PIKfive inhibitor, also inhibited viral replication, and therefore this appears to be a class effect. Human influenza is able to infect and replicate in a number of animal species used for pre-clinical evaluation (27) and most work of this nature is conducted in mice or in ferrets. The inbred mouse strains are characterised as “semi-permissive” for the replication of human respiratory viruses, and viral replication and associated symptoms were more marked in ferrets (28). Therefore, we used H1N1 PR8, which is relatively adapted to mice. In addition, as clinical trials of oral Apilimod demonstrated limited systemic exposure, intra-nasal treatment was used to achieve appropriate concentrations locally. As demonstrated in this study, topical administration of Apilimod to the lungs through nostrils produced profound inhibition of virus production and exhibited a dose-dependent steep inhibition curve in H1N1 infected BALB/c mice (Figure 3).

As the human respiratory virus is semi permissive to animals and the outcome of cancer cell lines is not translational, ALI cultured primary airway cells have been used (29, 30). The cellular layer consisted of ciliated cells and some goblet cells, and the system shows apical shedding of progeny virions that are subsequently spread by the coordinated motion of the beating cilia, so mimicking human influenza infection (31, 32). In fact, addition of influenza at a low level of infectious particles (0.01 MOI) apically to ALI nasal epithelium, resulted in a robust, persistent infection which generated amplified viral concentrations over seven days (Figure 2E, and unpublished data). In this ALI system, Apilimod displayed a concentration dependent inhibition of influenza replication determined by TCID_50_ assay following daily apical exposure (mimicking a topical/aerosol delivery to the respiratory tract) on Day 0 and Day 1 post-inoculation.

A highly desirable feature of inhaled medicines is extended longer duration of action, thereby ensuring that therapeutic activity is maintained throughout the dosing interval. The persistence of action of Apilimod was not fully investigated, but at least once daily treatment retains effective antiviral activity in both ALI cultured fully differentiated nasal epithelium *in vitro,* as well as in mice *in vivo* (Figure 2 and Figure 3C). The observed persistence of effect of Apilimod may be particularly valuable in the context of the potential use of Apilimod in prophylaxis.

Apilimod did not show strong antiviral effects against RSV A2 in African green monkey cell line, but it demonstrated marked antiviral effects *in vitro* ALI human nasal epithelium and also mouse *in vivo*. The african green monkey cell line might be not suitable to evaluate an PIKfyve inhibitor due to toxicity. From our results with human primary nasal epithelium, the antiviral effect of Apilimod was confirmed in influenza and RSV, which are negative-negative sense single-stranded RNA viruses. Even more importantly, Apilimod also showed the antiviral effects against positive strand RNA viruses, such as HRV and CoV. Particularly, the effects against seasonal CoV were marked. Apilimod was also reported to show very potent inhibitor against SARS-CoV-2 infection in Vero cells, and also confirmed in Calu3 and primary lung tissue (15) although Dittmer et al., found the antiviral effects of Apilimod to be weaker in Calu3 cells compared to Vero cells (33).

Thus, Apilimod demonstrated promising anti-viral potentials in this study, but there are some limitations for interpretation. Firstly, we have not tested other animal species. Mouse has some drawback, especially for influenza infection due to antiviral MX-1 protein mutation (34). Ferrets for influenza and cotton rats for RSV would be preferred systems to test the effects of Apilimod in future, although we already evaluated the antiviral effects against H1N1 and RSV using translational ALI human epithelium. Secondary, we only tested PR8 influenza strain, largely adapted to mouse, in ALI epithelium and mice. As we observed the efficacy of Apilimod against other human influenza isolates (Table 1), the animal and ALI studies with human influenza clinical isolates should be conducted in future. Thirdly, we need several donors from ALI epithelium, as donor-donor variation is not negligible in general in this primary cell system. In fact, previously, we conducted power calculation on RSV-infection system in ALI epithelium, and 4 different donors will be required to evaluate anti-viral effects (24, 35). However, to resolve the issue partially, we used 4-5 different donor pooled ALI-nasal epithelium in this study, provided by Epithelix. Fourthly, although an advantage of pan-viral agents targeting host cell factors would be a limited development of resistant mutants, we did not evaluate the emergence of resistant mutans in this system yet. At least we confirmed Apilimod did not induce resistant variants up to 8 passages in 229E coronavirus infected MRC5 human fibroblast cells where Nirmatrelvir induced mutants after 5 passages (data not shown). Finally, a PIKfyve inhibitor potentially inhibits virus antigen presentation due to inhibition of endosome systems (36), which potentially inhibiting antigen specific T cell activation *in vivo*. Therefore, therapeutic treatment of Apilimod after virus infection is established might block antiviral immune responses. We designed experiment for prophylaxis or early intervention, but an impact of therapeutic treatment should be explored in future.

In summary, we found Apilimod as a potential host -directed pan-viral small-molecule inhibitor against respiratory viruses including influenza and RSV in ALI human epithelium *in vitro* and *in vivo* mouse and also against seasonal CoV and HRV *in vitro*, which constitutes a promising candidate for the treatment of respiratory virus infection in humans via intranasal delivery.

## MATERIALS AND METHODS

### Materials

Apilimod base, Apilimod mesylate, YM201636, Oseltamivir phosphate and Oseltamivir carboxylate were purchased from MedChemExpress LLC (Monmouth Junction, NJ), Ribavirin were purchased (confirmed by Pharmidex).

### Cells and virus

MDCK (ATCC® CCL-34™) and Human larynx epithelial (HEp-2) cells (ATCC® CCL-23™) were purchased from the American Type Culture Collection (ATCC, Manassas, VA) and maintained in 10% foetal bovine serum (FBS) supplemented DMEM with phenol red (# 4190-094: Life Technologies Ltd, Paisley, UK) at 37°C/5% CO_2_. MucilAir™ pooled donor nasal epithelium was provided as 24-well plate sized inserts by Epithelix Sàrl (Geneva, Switzerland). Twice weekly, MucilAir™ inserts were transferred to a new 24-well plate containing 780µL of MucilAir™ culture medium (EP04MM), and once weekly the apical surface was washed once with 400µL PBS. MucilAir™ cultures were incubated at 37°C, 5% CO_2_. RSV A2 Strain (#0709161v, NCPV, Public Health England, Wiltshire, UK or ATCC® VR-1540™) and influenza PR8 (ATCC® VR-95™) were propagated in HEp-2 and MDCK cells, respectively for in house *in vitro* work.

### Anti-viral panel screening

The effects of Apilimod mesylate and YM201636 against a panel of viruses were evaluated at Institute for Antiviral Research in Utah State University. The assays were conducted as follows: Plaque reduction assay (5 days) in HEp-2 cells for RSV A (Long strain), CPE (6 days) in LLC-MK2 7.1 cells for Parainfluenza (PIV3 C243 strain), CPE (10 days) in MRC5 cells for Measles virus (Edmonton strain), CPE (5 days) in A549 cells for Influenza A virus (A/PR/34 strain), CPE (3 days) in HeLa cells for human Rhinovirus (HRV16 strain), CPE (5 days) in CEM-SS cells for HIV-1(IIIB strain), CPE (4 days) in Vero cells for Herpes Simplex Virus-1(HF strain), GT1b replicon luciferase assays (3 days) in Huh-7 cells for HCV. PC786 was applied 2 hrs before infection.

### Cytopathic effects (CPE) assay and cell viability (To be conducted)

For the assay of PR8 strains, 96 well format resazurin-based CPE assay was conducted. Approximately 24 hrs after the cells were seeded into 96 well black plates (200uL of 5% FBS phenol red-free DMEM/well) at a density of 3x10^4^ cells/ml, the cells were infected with PR8 (multiplicity of infection (MOI) of 0.2). Plain medium was also added to non-treatment wells as non-infection controls. Immediately after the infection, the compounds and neat DMSO (0.5µL /well) were added as appropriate to give a final concentration of 0.5% DMSO across all wells. The plates were incubated for 5 days (34°C, 5% CO_2_). After removing supernatant, 200µL of resazurin solution (0.0015% in PBS) was added to each well and the plates were incubated for a further 1 hr at 37°C/5% CO_2_. The fluorescence of each well [545 nm (excitation) / 590 nm (emission)] was then determined. The percentage inhibition for each well was calculated against infection control and the IC_50_ and IC_90_ values were calculated from the concentration-response curve generated for each test compound. For assessment of cell viability, test compounds or neat DMSO as vehicle (1µL) were added to each well of confluent MDCK culture in 96 well plates (200µL of 2.5% FBS DMEM/well), and incubated for 5 days (34°C, 5% CO_2_). Cells were then incubated with resazurin solution and the fluorescence level of resorufin (metabolized materials) was determined as described above. Where appropriate, a 50% cytotoxicity concentration (CC_50_) value was calculated from the concentration-response curve generated for each test compound.

### H1N1 or RSV infection and treatment on ALI cultured nasal epithelium

On the day of infection (‘Day 0’), the apical surface of each insert was washed once with 300 µL of sterile PBS and the inserts were then transferred to new 24-well plates containing 780 µL of fresh MucilAir™ culture medium (Epithelix Sarl. Switzerland). PR8 virus stock was diluted in MucilAir™ culture medium to give a final inoculation concentration of 4000 PFU in 100 µL (an approximate MOI of 0.02) for influenza, respectively, and incubated onto cells for one hour at 34°C, 5% CO_2_. Virus inoculum was removed with a pipette and inserts were washed twice with 300µL of sterile media. A Day 0 sample was harvested by adding 200 µL of culture media to the apical surface of each well for 10 minutes. The 200µL sample was then removed and transferred to 0.5ml tubes and the tubes were stored at -80°C. This harvesting procedure was repeated daily. Transepithelial electrical resistance (TEER) was measured to investigate the integrity of tight junction dynamics in air-liquid interface cultured pseudostratified epithelium before and after influenza infection as a surrogate for epithelial damage. Chopstick-electrodes were placed in the apical and basolateral chambers and the TEER was measured using a dedicated volt-ohm meter (EVOM2, Epithelial Volt/Ohm Meter for TEER) and expressed as Ohm/cm^2^. 50µL of compound solution was applied on Day 0, and incubated for 10min, and virus inoculum was applied on top of the treatment for 1hour. On Day1, 50µL of compound solution was applied, and incubated for 15min, and then removed. Oseltamivir was applied in basolateral chamber 10 min before virus inoculation, and it applied again on Day1. Viral load was assessed by TCID_50_ assay.

For the H1N1 study in Table 3 and RSV A2 study, 50µL of the compound solution or PBS was carefully applied to apical site and incubate at 37°C/5% CO_2_ for 15min. H1N1 PR8 virus or RSV A2 virus stock was diluted in MucilAir™ culture medium to give a final inoculation concentration of 4000 PFU in 50 µL, and applied to apical site on top of the treatment (2000 PFU in final, approximately an MOI of 0.01). Cells were incubated for another hour at 37°C/5% CO_2_, and then the apical media including virus inoculum was carefully removed with a pipette. After apical wash twice with PBS, the third apical wash was collected as Day 0 sample. On Day 1 (next day), 300µL of warm PBS was applied to the apical surface, and after 5 min, the first wash was collected for viral load. Following the first wash, 50µL of compound solution or PBS was applied to the apical surface and removed after 30min incubation. On Day 2, 300µL of warm PBS was applied to apical surface, and after 5 min, the first wash was collected daily for viral load assessment by plaque assay. Oseltamivir or Ribavirin was also applied to basal chamber for H1N1 and RSV A2 study, respectively.

### *In vivo* H1N1 infection

BALB/c mice (male, 20–30g) were inoculated intranasally on Day 0 with H1N1 PR8 (0.65 x 10^5^ PFU/mouse) in Pharmidex UK. Animals were sacrificed 3 days after the inoculation, and broncho alveolar lavage and nasal lavage were collected for inflammation. The lungs and nasal tissue were also collected for preparation of tissue homogenate. Apilimod base (Santa Cruz Biotechnolgy, Heidelberg, Germany) was prepared as suspension in 10% DMSO/90% isotonic saline and delivered using an intranasal installation of 40 μL/mouse at doses of 0.5, or 2 mg/mL on Day -1, again on Day 0 (1 hour before inoculation) and then once daily on Days 1, 2 and 3 post-inoculation. Apilimod mesylate was also prepared as solution in sterile water and treated intranasally. Oseltamivir phosphate (Sigma-Aldrich) was suspended in phosphate buffer saline, and was administered by oral gavage (10mL/kg). Viral load was determined by plaque assay using MDCK cells in the presence of L-1-tosylamido-2-phenylethyl chloromethyl ketone (TPCK) treated trypsin 1.0 µg/ml, and the plaque was detected by crystal violet staining 3 days after inoculation. The detection limit was 50 PFU/mL of tissue homogenate. Viral load was corrected to the weight of lung homogenate, and shown as PFU/g tissue. All work will be conducted under United Kingdom home office project licence PP0969406.

### *In vivo* RSV infection

BALB/c mice (male, 20–30g) were inoculated intranasally on Day 0 with RSV A2 (0.65 x 10^5^ PFU/mouse) in Pharmidex UK. Animals were sacrificed 4 days after the inoculation, and the lungs were collected for preparation of lung homogenate. Apilimod base was prepared as suspension in 10% DMSO/90% isotonic saline and delivered using an intratracheal installation of 20 μL/mouse or intranasal installation of 40 μL/mouse at doses of 0.2, or 2 mg/mL on Day -1, again on Day 0 (1 hour before inoculation) and then once daily on Days 1, 2 and 3 post-inoculations. All work will be conducted under United Kingdom home office project licence PP0969406.

### TCID_50_ assay for H1N1

Serial dilutions of the apical wash samples with media containing 0.1 μg/ml TPCK Trypsin were applied to plate wells containing an 80% confluency of MDCK cells and incubated at 35°C with 5% CO_2 for_ 3-5 days until the cytopathic effects of vehicle control became visible. TCID_50_ was calculated using the Reed-Muench formula (37).

### Plaque assay for H1N1

Serial dilutions of the apical wash samples from ALI culture or mouse lavage samples were applied to plate wells containing an 80% confluency of MDCK cells. The inoculated cells were incubated at 37°C for 1 h after which the inoculum was removed from the wells and the cells washed (twice) with PBS before applying an overlay of 1% methylcellulose agar media (inc. growth media plus TPCK trypsin) to each well. Once the agar media has set the plates was placed in an incubator at 37°C with 5% CO_2_ for 3 days after which the resulting plaques will counted. A second count was conducted once the agar has been removed by fixing and staining the cells with crystal violet. Data were presented as mean ± sem pfu for each group.

### Plaque assay for RSV

HEp2 cells was grown in 24-well plates prior to infection in DMEM containing 10% v/v FBS until they attain 100% confluency. Apical wash samples from ALI culture or mouse lavage were thawed out at room temperature and serial dilutions was prepared in serum-free DMEM. The growth medium from HEp2 cells were aspirated and replaced with 300 µL of serially diluted lung homogenate (along with stock RSV only positive control) and left to infect at 37°C/5% CO_2_ for four hours. The infectious media was then be aspirated and replaced with 500 µL Plaque Assay Overlay (1% w/v methylcellulose in MEM, 2% v/v FBS, 1% w/v pen/strep, 0.5 µg/ml amphotericin B), and left for 7 days at 37°C/5% CO_2_. Cells were fixed with ice-cold methanol for 10 minutes after which they were washed twice with sterile PBS. Anti-RSV F-protein antibody [2F7] was diluted to a 1:150 concentration in blocking buffer (5% w/v powdered milk (Marvel) in 0.05% v/v PBS-Tween 20) and 150 µL was added to cells for 2 hours at room temperature with shaking. Cells were washed 2x using PBS before 150 µl of secondary antibody (goat anti-mouse/HRP conjugate) diluted 1:400 in blocking buffer were added to cells for 1 hour at room temperature, with shaking. The secondary antibody solution was removed, and cells was washed twice with PBS. 150 µL of the metal-enhanced development substrate DAB prepared in ultra-pure water was applied to the cells until plaques are visible. Plaques was counted by eye and confirmed using light microscopy, allowing the calculation of plaque-forming units per mL.

### Lactose Dehydrogenase (LDH) Assay

LDH concentrations of collected samples were measured using commercial ELISA kit (Abcam, UK) as per the manufacturer’s instructions. Optical density was measured at 450 nM using a microplate reader (SpectraMax 340PC). Concentrations of LDH was determined using SoftMax Pro v. 6.4 (Molecular Devices).

### Statistical analysis

Results were represented as mean ± standard error of the mean. The IC_50,_ IC _90_ and CC_50_ values were calculated using GraphPad Prism (GraphPad Software Inc., La Jolla, CA). The safety index was calculated as the ratio of the CC_50_ and IC_50_ values. Multiple comparison was performed by ANOVA followed by Dunnett’s multiple comparison test performed using the PRISM 6^®^ software program. If no significance was achieved using ANOVA analysis, non-parametric Kruskal-Wallis analysis followed by Dunn’s multiple comparison test was conducted. The comparison between two groups was performed by unpaired *t*-test with Welch’s correction or Mann-Whitney test. Statistical significance was defined as *p*<0.05.

## ACKNOWLEDGEMENT

We thank Jag Shur, SubIntro Ltd. for insightful discussion.

K.I., G.R. J.B conceived and designed the experiments.

J.B., H.O., L.D., I.K. performed the experiments.

The first draft of the manuscript was written by K.Ito. All authors contributed to the technical interpretation, the interpretation of the result, and the editing of the manuscript.

## DATA AVAILABILITY STATEMENT

Data will be available on request.

## REFERENCES

1. Linden D, Guo-Parke H, Coyle PV, Fairley D, McAuley DF, Taggart CC, Kidney J. 2019. Respiratory viral infection: a potential "missing link" in the pathogenesis of COPD. Eur Respir Rev 28.

2. Zheng XY, Xu YJ, Guan WJ, Lin LF. 2018. Regional, age and respiratory-secretion-specific prevalence of respiratory viruses associated with asthma exacerbation: a literature review. Arch Virol 163:845–853.

3. Cookson W, Moffatt M, Rapeport G, Quint J. 2022. A Pandemic Lesson for Global Lung Diseases: Exacerbations Are Preventable. Am J Respir Crit Care Med 205:1271–1280.

4. Jin Y, Deng Z, Zhu T. 2022. Membrane protein trafficking in the anti-tumor immune response: work of endosomal-lysosomal system. Cancer Cell Int 22:413.

5. Yong X, Mao L, Shen X, Zhang Z, Billadeau DD, Jia D. 2021. Targeting Endosomal Recycling Pathways by Bacterial and Viral Pathogens. Front Cell Dev Biol 9:648024.

6. Nelson EA, Dyall J, Hoenen T, Barnes AB, Zhou H, Liang JY, Michelotti J, Dewey WH, DeWald LE, Bennett RS, Morris PJ, Guha R, Klumpp-Thomas C, McKnight C, Chen YC, Xu X, Wang A, Hughes E, Martin S, Thomas C, Jahrling PB, Hensley LE, Olinger GG, Jr., White JM. 2017. The phosphatidylinositol-3-phosphate 5-kinase inhibitor apilimod blocks filoviral entry and infection. PLoS Negl Trop Dis 11:e0005540.

7. Cai X, Xu Y, Cheung AK, Tomlinson RC, Alcazar-Roman A, Murphy L, Billich A, Zhang B, Feng Y, Klumpp M, Rondeau JM, Fazal AN, Wilson CJ, Myer V, Joberty G, Bouwmeester T, Labow MA, Finan PM, Porter JA, Ploegh HL, Baird D, De Camilli P, Tallarico JA, Huang Q. 2013. PIKfyve, a class III PI kinase, is the target of the small molecular IL-12/IL-23 inhibitor apilimod and a player in Toll-like receptor signaling. **Chem Biol** 20:912-921.

8. Wada Y, Cardinale I, Khatcherian A, Chu J, Kantor AB, Gottlieb AB, Tatsuta N, Jacobson E, Barsoum J, Krueger JG. 2012. Apilimod inhibits the production of IL-12 and IL-23 and reduces dendritic cell infiltration in psoriasis. PLoS One 7:e35069.

9. Krausz S, Boumans MJ, Gerlag DM, Lufkin J, van Kuijk AW, Bakker A, de Boer M, Lodde BM, Reedquist KA, Jacobson EW, O’Meara M, Tak PP. 2012. Brief report: a phase IIa, randomized, double-blind, placebo-controlled trial of apilimod mesylate, an interleukin-12/interleukin-23 inhibitor, in patients with rheumatoid arthritis. Arthritis Rheum 64:1750–1755.

10. Sands BE, Jacobson EW, Sylwestrowicz T, Younes Z, Dryden G, Fedorak R, Greenbloom S. 2010. Randomized, double-blind, placebo-controlled trial of the oral interleukin-12/23 inhibitor apilimod mesylate for treatment of active Crohn’s disease. Inflamm Bowel Dis 16:1209-1218.

11. Kang YL, Chou YY, Rothlauf PW, Liu Z, Soh TK, Cureton D, Case JB, Chen RE, Diamond MS, Whelan SPJ, Kirchhausen T. 2020. Inhibition of PIKfyve kinase prevents infection by Zaire ebolavirus and SARS-CoV-2. Proc Natl Acad Sci U S A 117:20803–20813.

12. Kumar P, Mathayan M, Smieszek SP, Przychodzen BP, Koprivica V, Birznieks G, Polymeropoulos MH, Prabhakar BS. 2022. Identification of potential COVID-19 treatment compounds which inhibit SARS Cov2 prototypic, Delta and Omicron variant infection. Virology 572:64–71.

13. Kettunen P, Lesnikova A, Rasanen N, Ojha R, Palmunen L, Laakso M, Lehtonen S, Kuusisto J, Pietilainen O, Saber SH, Joensuu M, Vapalahti OP, Koistinaho J, Rolova T, Balistreri G. 2023. SARS-CoV-2 Infection of Human Neurons Is TMPRSS2 Independent, Requires Endosomal Cell Entry, and Can Be Blocked by Inhibitors of Host Phosphoinositol-5 Kinase. J Virol doi:10.1128/jvi.00144-23:e0014423.

14. Luo Z, Liang Y, Tian M, Ruan Z, Su R, Shereen MA, Yin J, Wu K, Guo J, Zhang Q, Li Y, Wu J. 2023. Inhibition of PIKFYVE kinase interferes ESCRT pathway to suppress RNA virus replication. J Med Virol 95:e28527.

15. Riva L, Yuan S, Yin X, Martin-Sancho L, Matsunaga N, Burgstaller-Muehlbacher S, Pache L, De Jesus PP, Hull MV, Chang M, Chan JF, Cao J, Poon VK, Herbert K, Nguyen TT, Pu Y, Nguyen C, Rubanov A, Martinez-Sobrido L, Liu WC, Miorin L, White KM, Johnson JR, Benner C, Sun R, Schultz PG, Su A, Garcia-Sastre A, Chatterjee AK, Yuen KY, Chanda SK. 2020. A Large-scale Drug Repositioning Survey for SARS-CoV-2 Antivirals. bioRxiv doi:10.1101/2020.04.16.044016.

16. Zarkoob H, Allue-Guardia A, Chen YC, Jung O, Garcia-Vilanova A, Song MJ, Park JG, Oladunni F, Miller J, Tung YT, Kosik I, Schultz D, Yewdell J, Torrelles JB, Martinez-Sobrido L, Cherry S, Ferrer M, Lee EM. 2021. Modeling SARS-CoV-2 and Influenza Infections and Antiviral Treatments in Human Lung Epithelial Tissue Equivalents. bioRxiv doi:10.1101/2021.05.11.443693.

17. Baldassi D, Gabold B, Merkel O. 2021. Air-liquid interface cultures of the healthy and diseased human respiratory tract: promises, challenges and future directions. Adv Nanobiomed Res 1.

18. Michi AN, Proud D. 2021. A toolbox for studying respiratory viral infections using air-liquid interface cultures of human airway epithelial cells. Am J Physiol Lung Cell Mol Physiol 321:L263–L280.

19. Liu J, Cao R, Xu M, Wang X, Zhang H, Hu H, Li Y, Hu Z, Zhong W, Wang M. 2020. Hydroxychloroquine, a less toxic derivative of chloroquine, is effective in inhibiting SARS-CoV-2 infection in vitro. Cell Discov 6:16.

20. Cochin M, Touret F, Driouich JS, Moureau G, Petit PR, Laprie C, Solas C, de Lamballerie X, Nougairede A. 2022. Hydroxychloroquine and azithromycin used alone or combined are not effective against SARS-CoV-2 ex vivo and in a hamster model. Antiviral Res 197:105212.

21. Deng J, Zhou F, Heybati K, Ali S, Zuo QK, Hou W, Dhivagaran T, Ramaraju HB, Chang O, Wong CY, Silver Z. 2021. Efficacy of chloroquine and hydroxychloroquine for the treatment of hospitalized COVID-19 patients: a meta-analysis. Future Virol doi:10.2217/fvl-2021-0119.

22. Maisonnasse P, Guedj J, Contreras V, Behillil S, Solas C, Marlin R, Naninck T, Pizzorno A, Lemaitre J, Goncalves A, Kahlaoui N, Terrier O, Fang RHT, Enouf V, Dereuddre-Bosquet N, Brisebarre A, Touret F, Chapon C, Hoen B, Lina B, Calatrava MR, van der Werf S, de Lamballerie X, Le Grand R. 2020. Hydroxychloroquine use against SARS-CoV-2 infection in non-human primates. Nature 585:584–587.

23. Killingley B, Mann AJ, Kalinova M, Boyers A, Goonawardane N, Zhou J, Lindsell K, Hare SS, Brown J, Frise R, Smith E, Hopkins C, Noulin N, Londt B, Wilkinson T, Harden S, McShane H, Baillet M, Gilbert A, Jacobs M, Charman C, Mande P, Nguyen-Van-Tam JS, Semple MG, Read RC, Ferguson NM, Openshaw PJ, Rapeport G, Barclay WS, Catchpole AP, Chiu C. 2022. Safety, tolerability and viral kinetics during SARS-CoV-2 human challenge in young adults. Nat Med 28:1031–1041.

24. Brookes DW, Coates M, Allen H, Daly L, Constant S, Huang S, Hows M, Davis A, Cass L, Ayrton J, Knowles I, Strong P, Rapeport G, Ito K. 2018. Late therapeutic intervention with a respiratory syncytial virus L-protein polymerase inhibitor, PC786, on respiratory syncytial virus infection in human airway epithelium. Br J Pharmacol doi:10.1111/bph.14221.

25. DeVincenzo J, Cass L, Murray A, Woodward K, Meals E, Coates M, Daly L, Wheeler V, Mori J, Brindley C, Davis A, McCurdy M, Ito K, Murray B, Strong P, Rapeport G. 2022. Safety and Antiviral Effects of Nebulized PC786 in a Respiratory Syncytial Virus Challenge Study. J Infect Dis 225:2087–2096.

26. Han A, Czajkowski LM, Donaldson A, Baus HA, Reed SM, Athota RS, Bristol T, Rosas LA, Cervantes-Medina A, Taubenberger JK, Memoli MJ. 2019. A Dose-finding Study of a Wild-type Influenza A(H3N2) Virus in a Healthy Volunteer Human Challenge Model. Clin Infect Dis 69:2082–2090.

27. Bem RA, Domachowske JB, Rosenberg HF. 2011. Animal models of human respiratory syncytial virus disease. Am J Physiol Lung Cell Mol Physiol 301:L148–156.

28. Nguyen TQ, Rollon R, Choi YK. 2021. Animal Models for Influenza Research: Strengths and Weaknesses. Viruses 13.

29. Liu X, Wu Y, Rong L. 2020. Conditionally Reprogrammed Human Normal Airway Epithelial Cells at ALI: A Physiological Model for Emerging Viruses. Virol Sin 35:280–289.

30. Heinen N, Klohn M, Steinmann E, Pfaender S. 2021. In Vitro Lung Models and Their Application to Study SARS-CoV-2 Pathogenesis and Disease. Viruses 13.

31. Zhang L, Peeples ME, Boucher RC, Collins PL, Pickles RJ. 2002. Respiratory syncytial virus infection of human airway epithelial cells is polarized, specific to ciliated cells, and without obvious cytopathology. J Virol 76:5654–5666.

32. Villenave R, Shields MD, Power UF. 2013. Respiratory syncytial virus interaction with human airway epithelium. Trends Microbiol 21:238–244.

33. Dittmar M, Lee JS, Whig K, Segrist E, Li M, Kamalia B, Castellana L, Ayyanathan K, Cardenas-Diaz FL, Morrisey EE, Truitt R, Yang W, Jurado K, Samby K, Ramage H, Schultz DC, Cherry S. 2021. Drug repurposing screens reveal cell-type-specific entry pathways and FDA-approved drugs active against SARS-Cov-2. Cell Rep 35:108959.

34. Staeheli P, Grob R, Meier E, Sutcliffe JG, Haller O. 1988. Influenza virus-susceptible mice carry Mx genes with a large deletion or a nonsense mutation. Mol Cell Biol 8:4518–4523.

35. Ito K, Daly L, Coates M. 2023. An impact of age on respiratory syncytial virus infection in air-liquid-interface culture bronchial epithelium. Front Med (Lausanne) 10:1144050.

36. Baranov MV, Bianchi F, van den Bogaart G. 2020. The PIKfyve Inhibitor Apilimod: A Double-Edged Sword against COVID-19. Cells 10.

37. Bullen CK, Davis SL, Looney MM. 2022. Quantification of Infectious SARS-CoV-2 by the 50% Tissue Culture Infectious Dose Endpoint Dilution Assay. Methods Mol Biol 2452:131–146.

